# Bio-Accelerated Weathering of Ultramafic Minerals with *Gluconobacter oxydans*

**DOI:** 10.1101/2024.11.25.625253

**Authors:** Joseph J. Lee, Luke Plante, Brooke Pian, Sabrina Marecos, Sean A. Medin, Jacob D. Klug, Matthew C. Reid, Greeshma Gadikota, Esteban Gazel, Buz Barstow

## Abstract

Ultramafic rocks are an abundant source of cations for CO_2_ mineralization (*e.g*., Mg) and elements for sustainability technologies (*e.g*., Ni, Cr, Mn, Co, Al). However, there is no industrially useful process for dissolving ultramafic materials to release cations for CO_2_ sequestration or mining them for energy-critical elements. Weathering of ultramafic rocks by rainwater, release of metal cations, and subsequent CO_2_ mineralization already naturally sequesters CO_2_ from the atmosphere, but this natural process will take thousands to hundreds of thousands of years to remove excess anthropogenic CO_2_, far too late to deal with global warming that will happen over the next century. Mechanical acceleration of weathering by grinding can accelerate cation release but is prohibitively expensive. In this article we show that gluconic acid-based lixiviants produced by the mineral-dissolving microbe *Gluconobacter oxydans* accelerate leaching of Mg^2+^ by 20× over deionized water, and that leaching of Mg, Mn, Fe, Co, and Ni further improves by 73% from 24 to 96 hours. At low pulp density (1%) the *G. oxydans* biolixiviant is only 6% more effective than gluconic acid. But, at 60% pulp density the *G. oxydans* biolixiviant is 3.2× more effective than just gluconic acid. We demonstrate that biolixiviants made with cellulosic hydrolysate are not significantly worse than biolixiviants made with glucose, dramatically improving the feedstock available for bioleaching. Finally, we demonstrate that we can reduce the number of carbon atoms in the biolixiviant feedstock (*e.g.*, glucose or cellulosic hydrolysate) needed to release one Mg^2+^ ion and mineralize one atom of carbon from CO_2_ from 525 to 1.

## Introduction

The accumulation of over a trillion tonnes of anthropogenic CO_2_ in the Earth’s atmosphere^1–4^ driving global climate change is one of the most pressing challenges of our time. To address this, the IPCC’s Special Report in 2018 stresses the necessity of deploying carbon dioxide removal (CDR) technology capable of removing tens of gigatonnes of CO_2_ from the atmosphere annually, paired with carbon-neutral energy technologies to prevent global warming from exceeding 1.5°C above pre-industrial levels^5^.

Mineralization of CO_2_ is a highly attractive approach for permanent sequestration of CO_2_, as it converts atmospheric carbon into solid minerals as a permanent sink^6,7^. Carbon mineralization proceeds through a two-step process. First, ultramafic rocks are dissolved by weathering to release magnesium cations (**Note S1**). In the second step, magnesium ions are reacted with dissolved carbonate to form magnesite (MgCO_3_), a stable mineral that permanently stores CO_2_^8^ (**Note S1**). A schematic of a proposed indirect carbonation method is shown in **Figure 1**.

**Figure 1.**
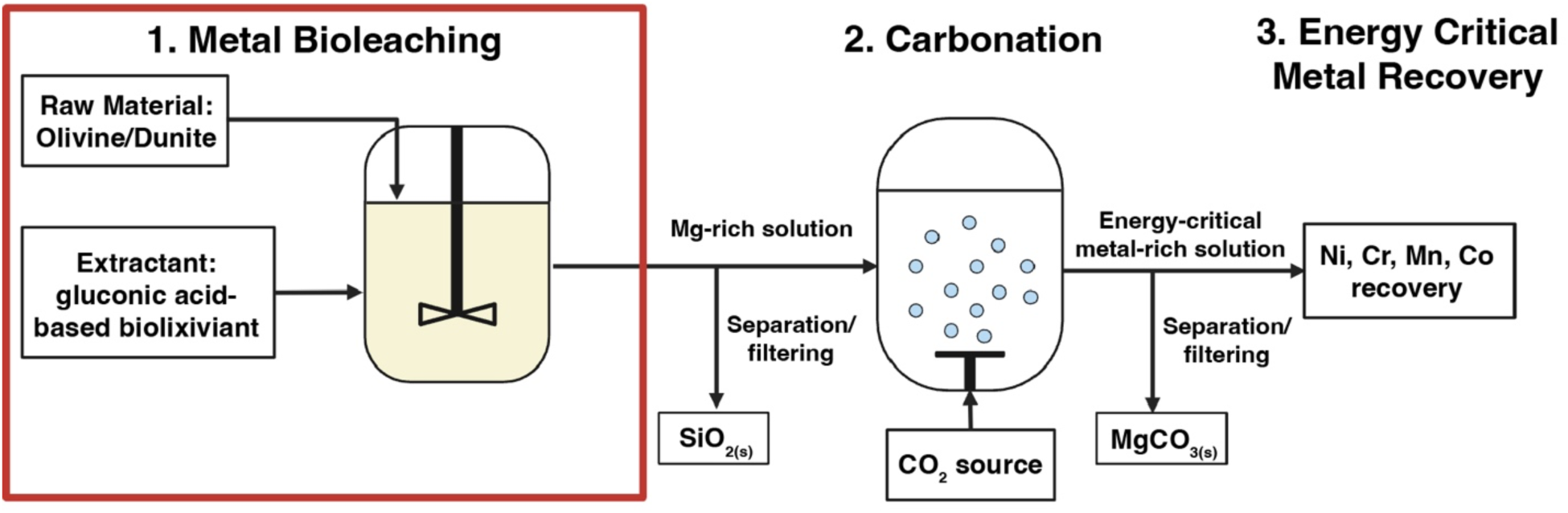
Proposed system for bio-accelerated weathering of ultramafic materials for carbon mineralization. This work focuses on the rate-limiting step, the metal leaching stage (enclosed in red box), where *G. oxydans* biolixiviant leaches magnesium ions and other metals from dunite. Downstream carbonation results in magnesite formation for CO_2_ storage, and metal recovery done chemically or electrochemically can extract energy-critical metals for commercial use.

Naturally occurring weathering of ultramafic rocks and subsequent carbon mineralization is thought to be responsible for removal of up to hundreds of millions of tonnes of CO_2_ each year^9–12^. In fact, because ultramafic rocks are so abundant^13,14^, reaction of all of the metal cations in all surface-accessible ultramafic material could capture 100 trillion tonnes of CO_2_ as carbonate minerals^15^ (at least 58× the excess CO_2_ in the atmosphere), making them a strong candidate for gigatonne-scale CDR^8,9,16^ However, natural carbon mineralization is partially offset by oxidation of rock organic carbon^12,17^. As a result, the time to naturally regulate atmospheric CO_2_ levels is estimated to be between tens and hundreds of thousands of years^9,12,18–20^

The large-scale deployment of carbon-neutral energy technologies will also be necessary to control climate change^5^, but it will require vast quantities of metals^21^. In addition to metal cations needed for CO_2_ sequestration, ultramafic minerals are a potential low-grade ore source for energy-critical metals such as nickel (batteries and stainless steel), chromium (concentrated solar and geothermal energy), manganese (steel and batteries), cobalt (batteries), and copper (wiring and wind turbines)^9,21–25^ (**Figure S1, Tables S1** and **S2**). This presents an opportunity to extract energy-critical metals from ultramafic rock while sequestering CO_2_.

However, extracting metals from ultramafic rocks poses a significant challenge. In aqueous mineral carbonation, the rate of carbon mineralization is limited by the slow kinetics of the leaching of metal ions from ultramafic sources^15,26,27^. While mechanical acceleration of ultramafic weathering through crushing, grinding, heating, chemically treating, and spreading over land is promising for its high sequestration potential and improved kinetics^9,16,28–30^, it requires large area footprints and high energy inputs involved in crushing, grinding, heating, and pressurizing, making it prohibitively expensive^31^. Likewise, technologies to extract the energy-critical elements present in low concentration in ultramafic rocks present formidable economic and environmental challenges^24,32,33^.

We hypothesize that microbially-accelerated mineral dissolution could overcome the slow kinetics and high costs^15^ of the dissolution of ultramafic material, releasing magnesium ions at a rate several orders of magnitude greater than in nature^34,35^. Over the past two decades, microbial mineral-dissolution has revolutionized mining copper and gold from low-grade ores—a process known as biomining. Bacteria such as *Acidithiobacillus ferrooxidans* and related species are responsible for producing about 15% of the world’s copper supply and about 5% of its gold supply^36^ through iron-redox-mediated bioleaching, a process in which microbes alter the redox state of metals to extract them from mineral substrates^37,38^. Significant research has been performed to improve industrial-scale Cu bioleaching with consortia including *A. ferrooxidans* and non-domesticated microbes, resulting in processes that are fairly mature. While copper biomining is the only industrialized biomining process, organisms and processes are in development for rare earth elements (REE)^39–44^ and lithium^45,46^.

Biolixiviants produced by *Gluconobacter oxydans* are a strong candidate for a sustainable and strong catalyst for CO_2_ mineralization using ultramafic rocks^47^. *G. oxydans* efficiently produces a gluconic acid-based mineral-dissolving solution, called a biolixiviant, when glucose is present^47,48^. Previous studies have shown that biolixiviants from *G. oxydans* can be more effective than just gluconic acid at leaching REE^42,49^. In a recent article our team identified genes that control the effectiveness of the *G. oxydans* biolixiviant at dissolving neodymium phosphate (synthetic monazite) but do not notably change its pH^50^. These results suggest that biologically-produced lixiviants could contain additional components, such as metal chelators, redox molecules, and flavins that improve overall metal leaching at fixed pH, allowing for more efficient and low-resource metal leaching. This could avoid the use of strong acids that can have harmful environmental effects such as acid mine drainage^51^.

Second, it is suspected that microbes interact with ultramafic materials in the wild, accelerating weathering processes^52–55^. Furthermore, laboratory mixed cultures have been shown to dissolve ultramafic material^56^. Meanwhile, in two companion articles, we demonstrate that three easily culturable isolated species (*G. oxydans*, *Penicillium simplicissimum* and *Sphingomonas desiccabilis*) can all accelerate the dissolution of ultramafic material^12^ and characterize mineral phase transformations due to bioleaching by *G. oxydans*^57^.

However, there has been little systematic study of microbial dissolution of ultramafic minerals. Furthermore, it remains unclear under what conditions a biolixiviant could be more effective than just an organic acid and why; could genetic engineering improve ultramafic bioleaching; what impact process parameters have on biological dissolution of ultramafic materials; the extraction efficiencies for energy critical elements; utilization of low-cost feedstock; and the carbon source demand of large-scale ultramafic bioleaching. To address these knowledge gaps, we provide some of the first systematic observations of dissolution of ultramafic minerals by the mineral-dissolving microbe, *G. oxydans* B58.

## Results and Discussion

In this work, we perform a study of *G. oxydans* bioleaching capability on dunite, an ultramafic rock composed of greater than 90% olivine (see composition in **Figure S1**, and **Tables S1** and **S2**). For accelerated weathering technologies, olivine-rich dunite is the preferred feedstocks due to olivine having the highest CO_2_ sequestration potential by mass, excellent carbonation kinetics, and is high abundance^58,59^.

This study benchmarks the bioleaching efficacy of glucose-based and lignocellulosic sugar-based biolixiviants produced by *G. oxydans* on dunite, comparing conditions of whole-cell and filtered (cell-free) biolixiviants, varying pulp densities, aerobic and anaerobic conditions, and short-term and long-term leaching. This study also investigates how a high-performing mutant strain of *G. oxydans* could improve leaching in ultramafic minerals, serving as a preliminary study of how genetic engineering could further improve bioleaching for carbon mineralization. We demonstrate that the effectiveness advantage of the *G. oxydans* biolixiviant over gluconic acid alone grows with increasing pulp density. Finally, we demonstrate that the number of carbon atoms that need to be supplied by sugar feedstocks can be reduced to match the number of carbon atoms potentially sequestered as a mineral.

### *G. oxydans* Biolixiviants Accelerate Magnesium and Energy-Critical Metal Leaching by 20× Over Deionized Water After 24 Hours

Biolixiviants from wild-type *Gluconobacter oxydans* effectively leached magnesium from ultramafic rocks (**Figure 2A**). The efficiency of extraction measures how much of the available metal is leached from the mineral, and is calculated from the ratio of metal concentrations in the leachate and the original concentration in the source multiplied by the pulp density (ratio of solid mass to liquid volume [g/L]),

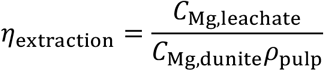

**Figure 2.**
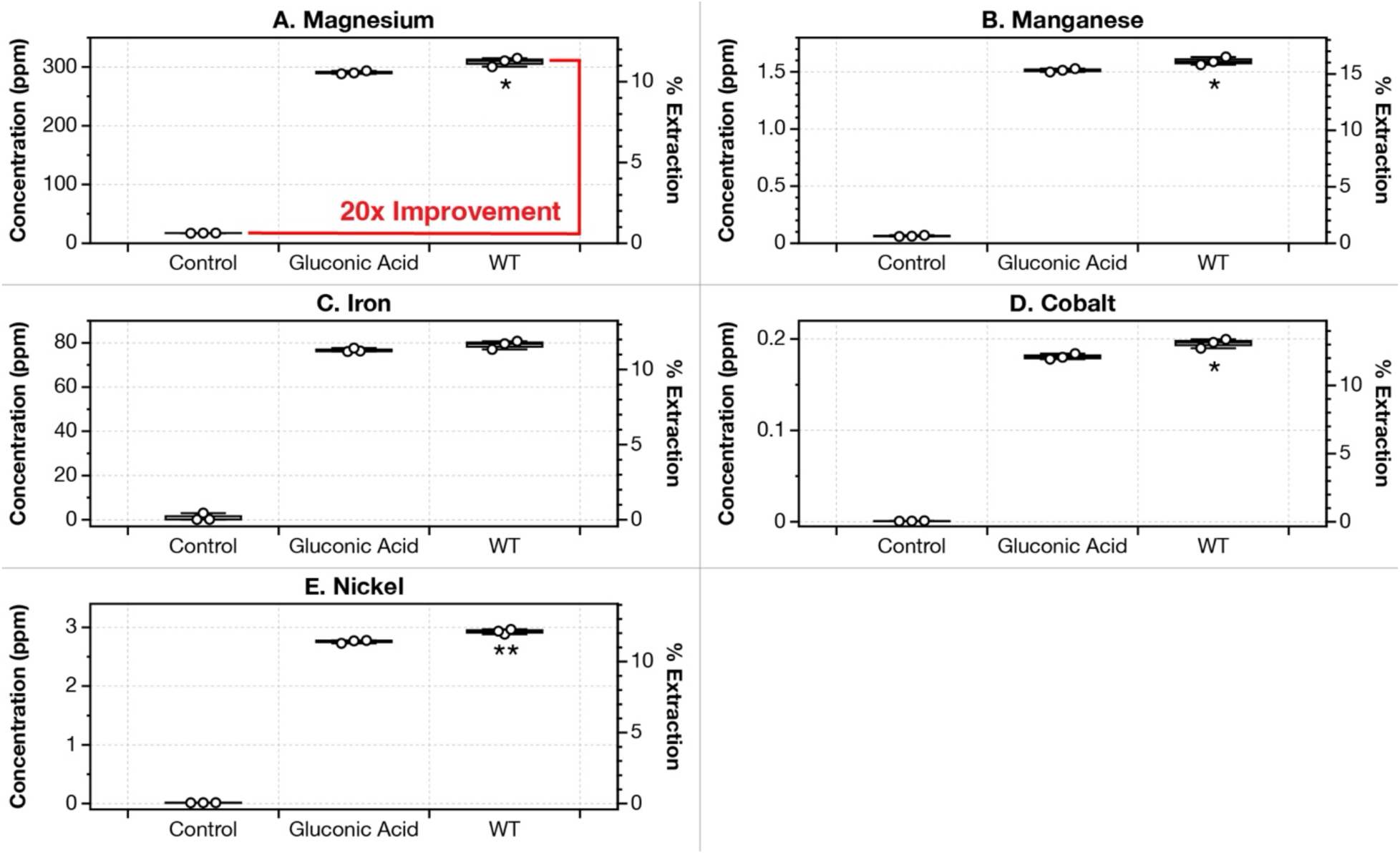
Leaching dunite at 1% pulp density with *G. oxydans* biolixiviant improves leaching of magnesium by 20-fold when compared to leaching with deionized water (control). However, the *G. oxydans* biolixiviant is only 6% better than just gluconic acid at the same pH. Bioleaching of dunite at 1% pulp density shaking at 22 °C for 24 hours was performed with wild-type *G. oxydans* biolixiviant, gluconic acid diluted to the same pH with deionized water, and a deionized water control. Panels **A** to **E** show leaching concentrations and extraction efficiencies of Mg, Mn, Fe, Co, and Ni. Wild-type *G. oxydans* biolixiviants slightly improved metal leaching compared to gluconic acid across all metals. Stars denote significant difference between gluconic acid leaching and wild-type biolixiviant leaching by a Welch’s two sample *t*-test, *p* < 0.05 (*), *p* < 0.01 (**), *p* < 0.001 (***).

Deionized water is used as a control for these experiments since it estimates the improvements made by bio-accelerated leaching to leaching from environmental water sources, such as rainwater. In traditional accelerated carbon mineralization with ultramafic rocks, crushed rocks are dispersed over land, where water from the environment is the solvent for magnesium^9,29,30^. Comparing the concentrations of magnesium between leaching with biolixiviants and water provides a rough estimate for the order of magnitude in which carbon mineralization is accelerated by *G. oxydans*.

After 24 hours of leaching with dunite at 1% pulp density, magnesium was leached with an extraction efficiency of 11% (300 ppm), corresponding to an increase of 20× when compared to the deionized water control (**Figure 2A**).

### Bioleaching of Metals by *G. oxydans* Increases by 73% From 24 to 96 Hours

The most efficiently extracted metal at 24 hours was Mn (**Figure 2B**), extracting over 15% of Mn from dunite (corresponding to a leachate concentration of 1.6 ppm). Fe, Co, and Ni were also leached highly, with efficiencies between 11% and 15% (**Figures 2C**, **D**, and **E**, respectively). Al and Cr were also leached, but much less efficiently (**Figure S2**). Cr was the least efficiently extracted metal, with extraction of no more than 0.3% under any condition (**Figure S2** right column).

When leaching for longer than 24 hours, bioleaching efficiency continues to increase (**Figure 3**). In a 96-hour bioleaching analysis of dunite at 1% pulp density, bioleaching improved between 24 and 96 hours by an average of 1.73-fold (or 73%) for Mg, Mn, Fe, Co, and Ni (**Figure 3**). After 96 hours, the leaching of magnesium in biolixiviant is 42 times that of water (**Figure 3**). This shows that biolixiviants from *G. oxydans* improve the overall leaching of metals at a rate faster than water.

**Figure 3.**
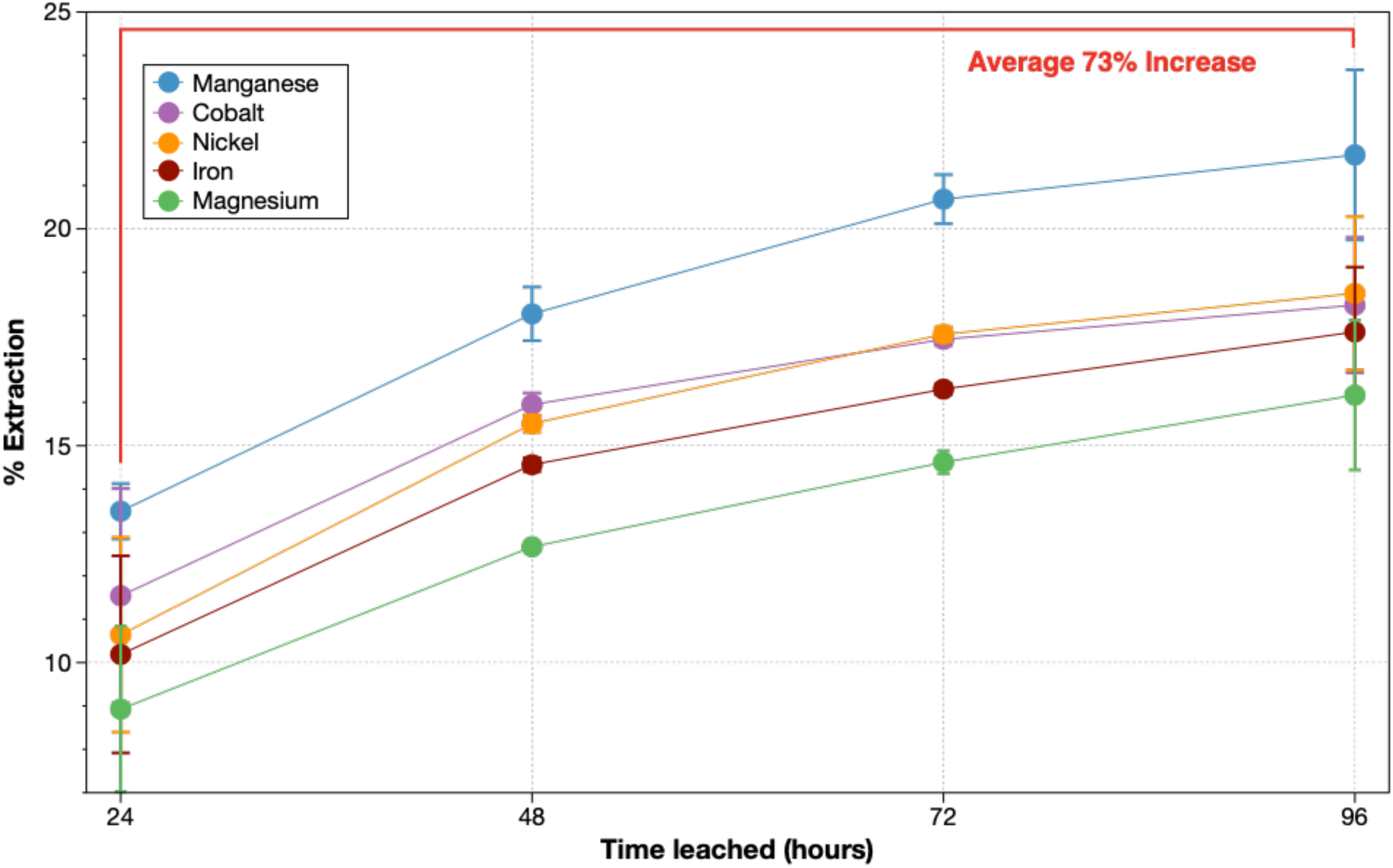
Increasing leaching time to 96 hours increases extraction efficiency by an average of 73% between 24 and 96 hours for Mg, Mn, Fe, Co, and Ni. Biolixiviants from wild-type *G. oxydans* leached dunite at 1% pulp density for 96 hours, shaking at 22°C. Mg, Mn, Fe, Co, and Ni concentrations were measured from leachates every 24 hours for 96 hours. Error bars indicate standard deviation of measurements from duplicate measurements.

### The *G. oxydans* Biolixiviant is Only 6% Better than Gluconic Acid at 1% Pulp Density

A secondary control was performed using only gluconic acid diluted with deionized water to the same pH as the acid biolixiviants produced by *G. oxydans*. Reed *et al*.^42^ and Antonick *et al*.^49^ demonstrated that biolixiviants produced by *G. oxydans* were noticeably more effective at bioleaching than gluconic acid alone. However, recent anecdotal evidence suggests this is not the case under all conditions. In our hands, when bioleaching ultramafic materials at low pulp density (1%), *G. oxydans*-produced biolixiviants only slightly outperform gluconic acid in leaching most metals (**Figure 2**). Wild-type *G. oxydans* biolixiviants leached only 6% more magnesium compared to pure gluconic acid that was diluted with ultrapure water to the same pH after 24 hours of leaching. Furthermore, at least with low pulp density leaching, filtering cells from the biolixiviant did not significantly affect bioleaching, at least under the conditions of these tests (**Figure 4**). This was done to assess if there was a significant residual effect of microbial activity on leaching after biolixiviant production. Furthermore, stoppering flasks during leaching to prevent gas exchange did not significantly affect bioleaching (**Figure S3**). Later in the article, we ask if there are conditions under which the *G. oxydans* biolixiviant has a significant performance advantage for bioleaching ultramafic materials.

**Figure 4.**
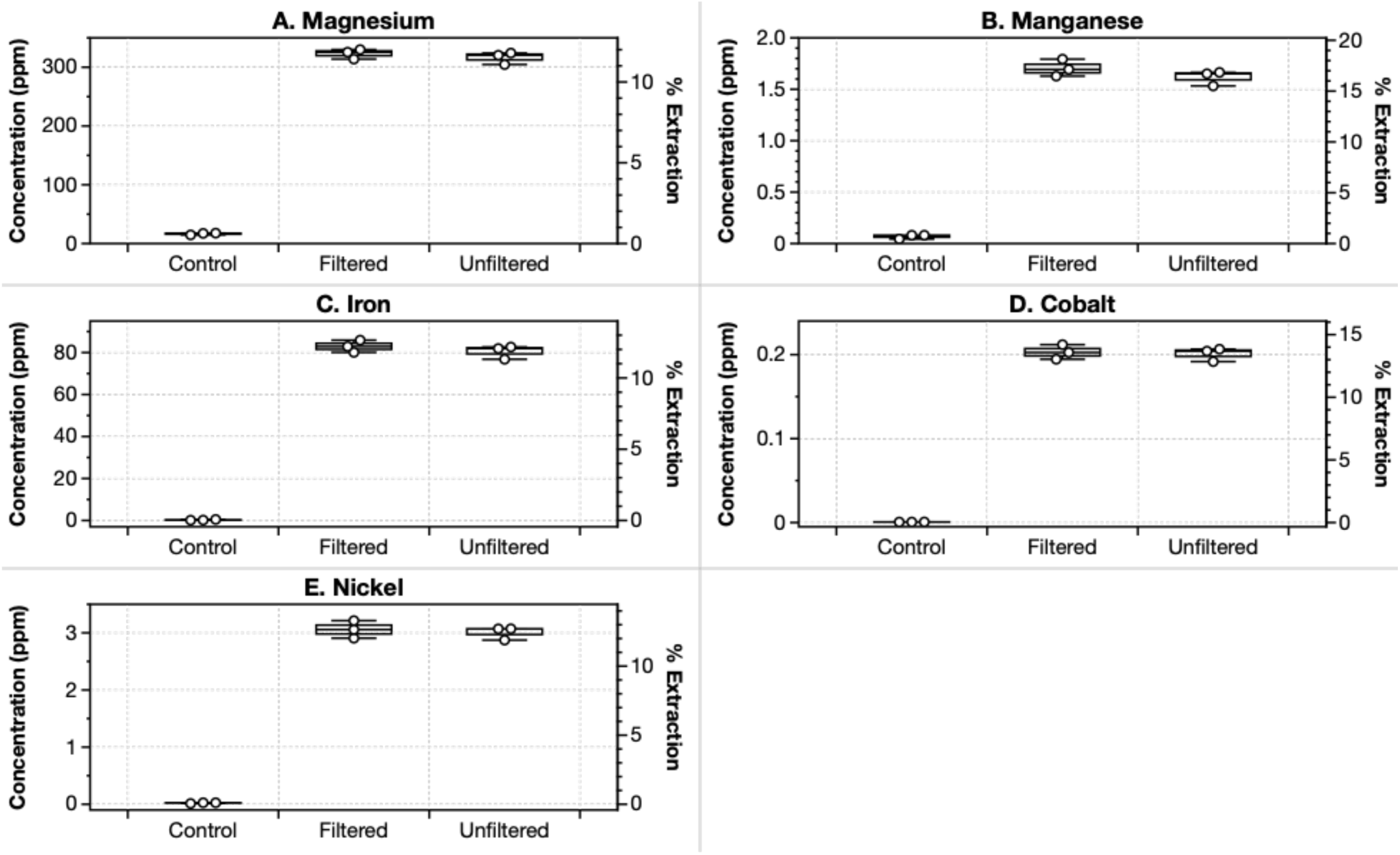
Filtering cells from the *G. oxydans* biolixiviant does not notably reduce the bioleaching of metals from dunite after 24-hour leaching. We compared Bioleaching of 1% dunite by sugar water (col. 1), with whole cell lixiviant (col. 2), and with a biolixiviant that was clarified by filtration with a 0.22 μm syringe filter (col. 3). Filtered and unfiltered biolixiviants were at identical pH, produced from duplicate colonies, and were added to ultramafic minerals in identical conditions. Filtering biolixiviants before bioleaching resulted in similar metal concentrations when compared to whole-cell biolixiviants when leaching dunite. A Welch’s two-sample t-test shows that no significant difference was detected between the unfiltered and filtered groups, with *p*-values ranging from 0.2 to 0.9. This indicates that based on a 24-hour leaching study, the presence of *G. oxydans* does not significantly improve or diminish the effect of the biolixiviant on leaching in short term leaching.

### Cellulosic Hydrolysate Can Be Used as an Alternative Sugar Feedstock for Bioleaching

Techno-economic analysis shows that glucose feedstock for biolixiviant production is the biggest cost driver in REE biomining with *G. oxydans*, accounting for almost half of the total cost^60^. Furthermore, the sheer scale that carbon sequestration needs to operate at, means that enormous amounts of feedstock will be required, potentially placing a significant demand on the world’s biomass supply^61^.

Lignocellulose, found in agricultural waste, can be a renewable carbon source for microbial growth and production once monomeric sugars are released through acid pretreatment and enzymatic hydrolysis, making a glucose-rich cellulosic hydrolysate^62,63^. Lignocellulosic sugars are being increasingly investigated as a potential alternative sugar source for several applications, including biofuels/bioethanol production and substrates for cellular production. Utilizing lignocellulose-derived sugars could serve as a high glucose replacement feedstock that would greatly improve the cost-efficiency of bioleaching for carbon capture technologies^64^. *G. oxydans* has been shown to be able to convert the glucose and xylose sugars in cellulosic hydrolysate into gluconic acid and xylonic acid at extremely high efficiencies^65–67^. Replacing pure glucose with cellulosic hydrolysate could further improve the economic and environmental viability of bio-accelerated weathering.

*G. oxydans* has been shown to efficiently produce gluconic acid and xylonic acid from lignocellulosic hydrolysate feedstocks^65–67^, making industrial-scale, bio-accelerated carbon mineralization far more economically feasible. We investigated if biolixiviants produced from cellulosic hydrolysate, rather than pure glucose, could be effective at leaching dunite. The cellulosic hydrolysate (CH) used in this study is a glucose-rich solution produced from the enzymatic hydrolysis of concentrated corn stover (**Table S3**).

Biolixiviants produced from cellulosic hydrolysate were on average 28% less effective in leaching magnesium compared to glucose-based biolixiviants (**Figure 5**). Leaching of Mn, Fe, Co, and Ni was also reduced by an average of 52% compared to glucose-based biolixiviants (**Figure 5**). Dunite was added at 1% pulp density, and a leaching comparison between CH-based and glucose-based lixiviants was conducted in parallel and in identical conditions. The cellulosic hydrolysate added was diluted with deionized water so that the total sugar concentrations were equal to the amount of glucose added for glucose-based biolixiviants.

**Figure 5.**
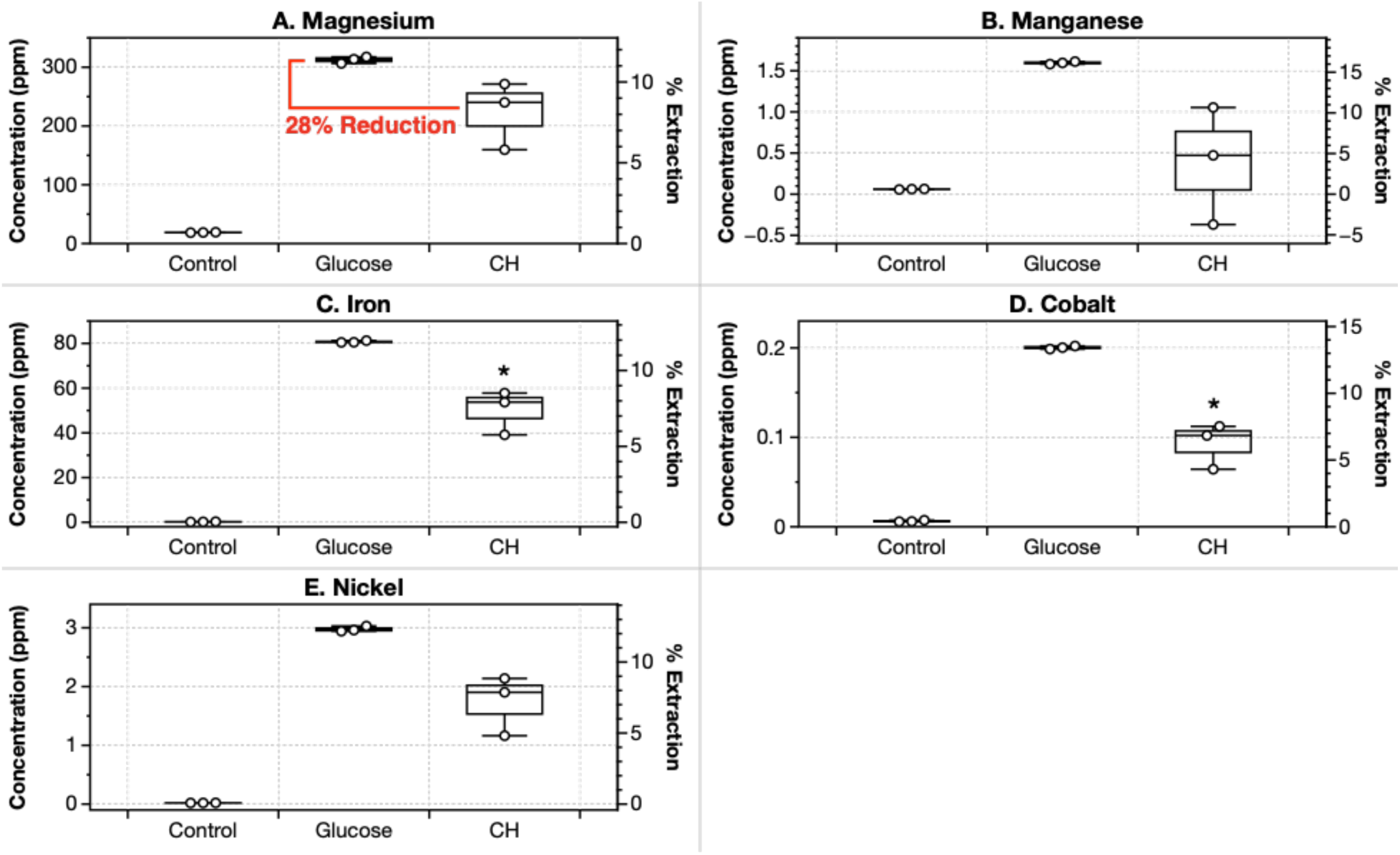
Use of cellulosic hydrolysate as a biolixiviant feedstock does not considerably reduce bioleaching efficiency. Panels **A** to **E** show comparison of bioleaching of Mg, Mn, Fe, Co, and Ni by *Gluconobacter oxydans* when using glucose and cellulosic hydrolysate (CH) as as biolixiviant feedstock. CH-based biolixiviant leaches magnesium on average at 72% the efficacy of glucose-based biolixiviant. Leaching was performed with 1% dunite shaking at 22°C for 24 hours. Stars denote significant difference compared with glucose-based biolixiviants by a Welch’s two sample *t*-test, *p* < 0.05 (*), *p* < 0.01 (**), *p* < 0.001 (***).

Despite a 28% average reduction in Mg leaching, we were unable to detect a statistically meaningful difference between glucose- and CH-based leaching of Mg using a Welch’s two-sample *t*-test. This was due to the small sample size and variability in CH-based biolixiviant leachate measurements. Data collected for CH-based biolixiviants resulted in greater variability than data with glucose-based biolixiviants. ICP-MS analysis of cellulosic hydrolysate samples showed that the CH used in this study contains trace concentrations of metals, possibly as a result of the manufacturing process (**Table S4**). When calculating the amounts of metal leached from CH-based biolixiviants, the concentrations of metal in a cellulosic hydrolysate control study were subtracted from the total detected to eliminate metal concentrations that were not extracted from dunite. In low pulp density studies, the concentrations of metals in the cellulosic hydrolysate feedstock interfered with metals leached, resulting in higher overall error.

Cellulosic hydrolysate produced from corn stover contains several toxins such as organic acids, furan derivatives, and phenolic compounds that could inhibit the growth of *G. oxydans* and its production of biolixiviant^68^. Cellulosic hydrolysate also significantly increases viscosity, resulting in lower oxygen transfer rates that limit the fermentation of glucose to gluconic acid^66^. Despite this, leaching results show that *G. oxydans* can produce a biolixiviant from cellulosic hydrolysate capable of leaching metals at concentrations significantly greater than controls; however, further analysis and experimentation are needed to show if CH can be a viable replacement to glucose without a significant reduction in performance.

Although cellulosic hydrolysate contains mild antibiotic properties due to its acidity and certain toxic compounds, this could be advantageous for the industrial scale use of it as a feedstock for biolixiviant production. *G. oxydans* has high acid resistance and is capable of growing and producing biolixiviant with cellulosic hydrolysate, while many other microbes have low tolerance mechanisms for it. This could allow for cellulosic hydrolysate to be used as an ultra-low-cost feedstock while also preventing the growth of competing microbes. The toxic components of cellulosic hydrolysate could also be a genetic engineering target for *G. oxydans*, where mutants that have better resistance mechanisms can be selected for to improve biolixiviant production.

### *G. oxydans* Mutant Strain Δ*pstS*, P112:*mgdh* Improved Magnesium Leaching by 12% Over Wild-Type

A preliminary study using the mutant strain *G. oxydans* Δ*pstS*, P_112_:*mgdh*^43^ shows that under identical growth and leaching conditions, mutant strains of *G. oxydans* can be more effective than wild-type in leaching magnesium and energy-critical metals from dunite.

A bioleaching analysis over 96 hours using dunite at 1% pulp density was conducted with biolixiviant from wild-type and mutant *G. oxydans*. Bioleachate samples were taken and analyzed every 24 hours up to 96 hours using ICP-MS. The strain used is *G. oxydans* Δ*pstS*, P_112_:*mgdh*, which was engineered to improve acid production for metal leaching^43^. It contains a clean deletion of the *pstS* gene which is involved in phosphate signaling and transport^44^, and an up-regulation with promoter P_112_ of the *mgdh* gene (Membrane-bound Glucose DeHydrogenase) which is critical for converting glucose into gluconic acid^69^. The Δ*pstS*, P_112_:*mgdh* mutant was previously shown to decrease the pH of biolixiviants compared to wild-type, and it increases the overall leaching of REE^43^.

The analysis shows that Δ*pstS*, P_112_:*mgdh* mutant biolixiviant increases magnesium leaching by an average of 12% over all samples taken between 24 and 96 hours of leaching. Mn, Fe, Co, and Ni had an average improvement in leaching of 12% using mutant biolixiviant compared to wild-type (**Figure 6**).

**Figure 6.**
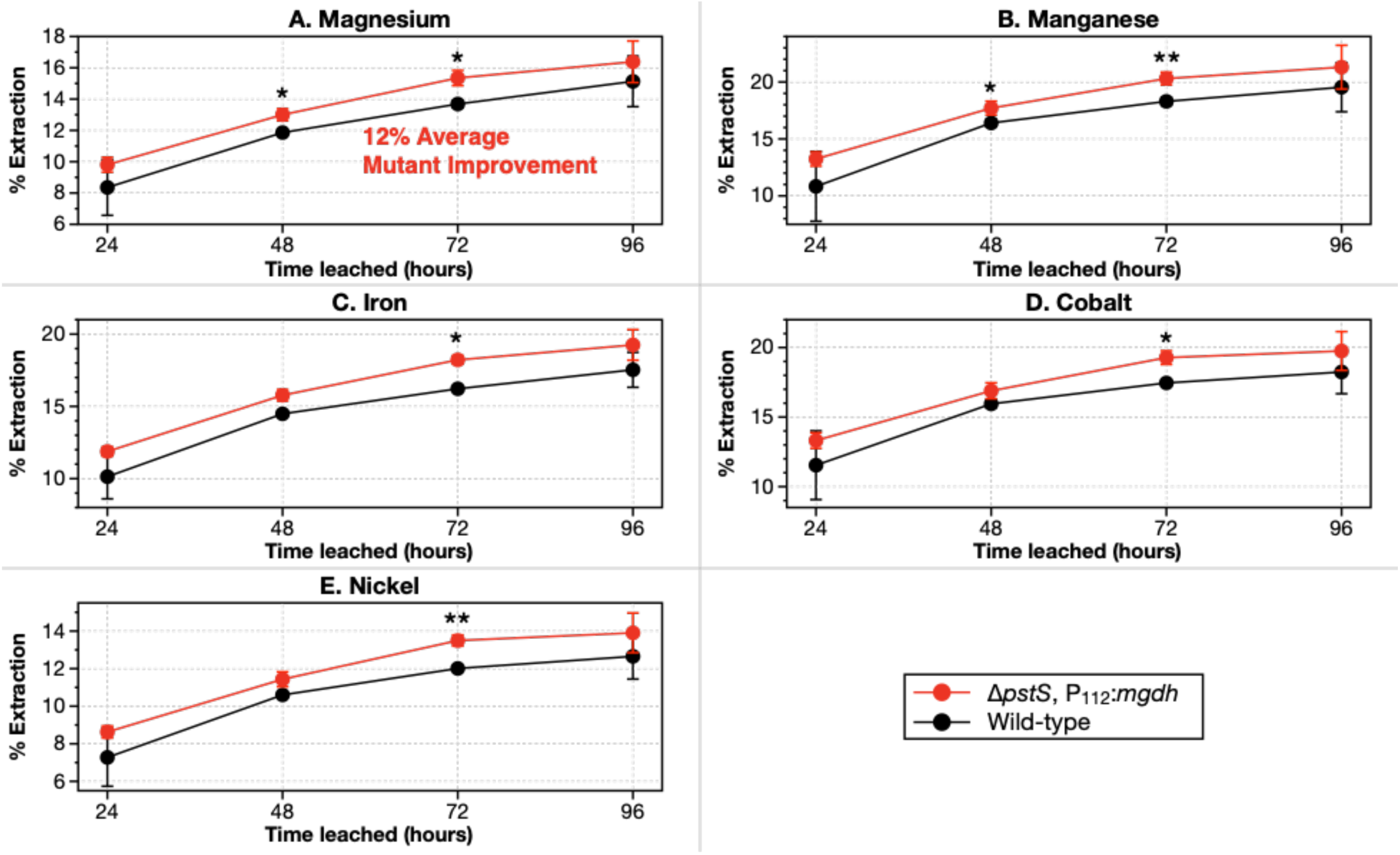
Bioleaching efficiency of magnesium increased by an average of 12% when leaching with a high performing *G. oxydans* mutant. The *G. oxydans* mutant contains a deletion of *pstS* phosphate signaling and transport gene, alongside up-regulation of the *mgdh* gene that codes for the Membrane-bound Glucose DeHydrogenase (MGDH) with the P_112_ promoter (*G. oxydans* Δ*pstS*, P_112_:*mdgh*). Leaching was done with dunite at 1% pulp density shaking at 22°C over 96 hours. Stars and dots denote a statistically significant difference between wild-type *G. oxydans* and *G. oxydans* Δ*pstS*, P_112_:*mdgh* mutant by a Welch’s two sample *t*-test, where, *p* < 0.1 (.), *p* < 0.05 (*), *p* < 0.01 (**), *p* < 0.001 (***).

The entire genome of *G. oxydans* is yet to be completely characterized, and several other genes may exist that are critical in controlling metal leaching from ultramafic rocks. Future work to characterize the genome of *G. oxydans* in terms of its functions related to ultramafic leaching is vital to engineer better mutant strains that can improve the performance of bio-accelerated weathering. The Δ*pstS*, P_112_:*mgdh* mutant is engineered to maximize gluconic acid production; however, studies have shown that for rare earths leaching, *G. oxydans* mutants can be screened to identify genes that specifically affect rare earth extraction^50^. Our research group has created a quality-controlled (QC) whole genome knockout collection for *G. oxydans* B58, containing a collection of 2,733 transposon disruption mutants, each with a different nonessential gene disrupted. This QC collection enables a high-throughput screen of the *G. oxydans* genome, selecting for significant changes to dunite leaching efficacy. This will provide us with a roadmap for genetically engineering *G. oxydans* for greater ultramafic bio-accelerated weathering capability.

### *G. oxydans* Biolixiviants are 3.2× More Effective than Gluconic Acid at 60% Pulp Density

At higher pulp densities and longer leaching times, *Gluconobacter oxydans* biolixiviants significantly outperformed gluconic acid at the same pH. We compared *Gluconobacter oxydans* biolixiviants and gluconic acid metal leaching with 5%, 10%, and 60% dunite pulp density, where leachate metal concentrations were measured every 24 hours for 72 hours. The gluconic acid was diluted with ultrapure deionized water to match the pH of biolixiviants, and leaching was performed in identical conditions with the same mineral substrates (DUN-2). After 72 hours of leaching, wild-type biolixiviants leached 1.74 times more magnesium than gluconic acid from 5% dunite, 2.87 times more from 10% dunite, and 3.25 times more from 60% dunite (**Figure 7**). At higher pulp densities, *Gluconobacter oxydans* biolixiviants have a greater improvement over gluconic acid. This analysis suggests that metal leaching from dunite is not only controlled by acidolysis, but might contain other biologically-synthesized components that amplify extraction^50^.

**Figure 7.**
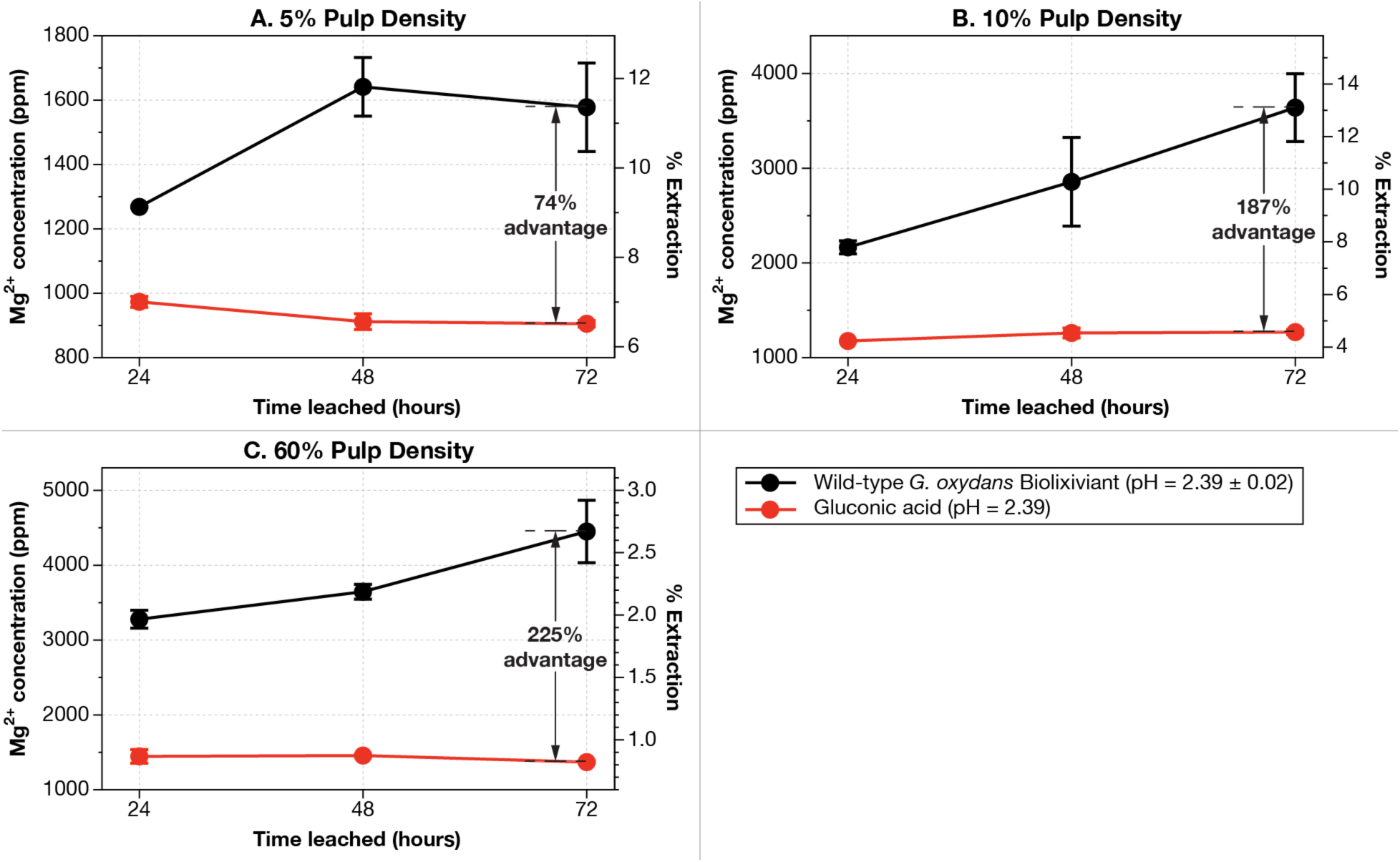
At 60% pulp density, the wild-type *G. oxydans* biolixiviant was 3.2× more effective at bioleaching dunite than gluconic acid diluted to the same pH with pure water. A comparative bioleaching study was conducted comparing the leaching efficacy of wild-type *G. oxydans* biolixiviants and just gluconic acid at the same pH. Magnesium concentrations in leachate were measured after 24, 48, and 72 hours of leaching with 5% (**A**), 10% (**B**), and 60% (**C**) dunite (DUN-2). At higher dunite pulp densities, the improvement of *Gluconobacter oxydans* biolixiviants compared to gluconic acid was amplified. After 72 hours, biolixiviants were 74% better than gluconic acid at leaching 5% dunite, 1.87 times better at leaching 10% dunite, and 2.25 times better at leaching 60% dunite. Stars denote significant difference between gluconic acid leaching and wild-type biolixiviant leaching by a Welch’s two sample *t*-test, *p* < 0.05 (*), *p* < 0.01 (**), *p* < 0.001 (***).

### Magnesium Leaching Relative to Glucose Input Can Be Greatly Improved by Process and Genetic Engineering

Glucose is the cost-limiting factor in utilizing *G. oxydans* biolixiviant as a weathering accelerant^60^. Furthermore, the sheer scale that carbon sequestration must operate in requires enormous amounts of microbial feedstock, possibly enough to make a significant demand on the world’s biomass supply^61^. Therefore, it is essential to prioritize maximizing carbon dioxide sequestration while minimizing feedstock consumption in order to mitigate the strain on worldwide biomass resources.

We define the carbon investment to carbon sequestration ratio, *k*_seq_, as the amount of carbon atoms in the feedstock (*e.g.*, glucose) needed to liberate one magnesium ion (and hence sequester the single atom of carbon in a CO_2_ molecule^12,61^),

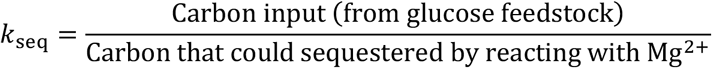

Lower *k*_seq_ values indicate that more magnesium was leached given a certain glucose input, while high *k*_seq_ values indicate relatively lower magnesium leaching for the same amount of feedstock consumed. A *k*_seq_ value of 6 for a glucose feedstock means that one glucose molecule is necessary to liberate one magnesium ion (and sequester one CO_2_). A *k*_seq_ value of 1 means that only one feedstock carbon atom is needed to eventually sequester one CO_2_ molecule.

We can use *k*_seq_ to estimate the price of carbon sequestration and the total amount of feedstock needed to sequester a useful amount of CO_2_ (*e.g.*, 20 gigatonnes of CO_2_ per year). The mass of feedstock, *M*_feedstock_, needed to sequester a given mass of CO_2_, 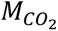,

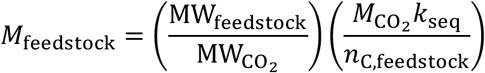

where, MW_feedstock_ is the molecular weight of the feedstock; 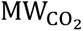 is the molecular weight of CO_2_, and *n*_C,feedstock_ is the average number of carbon atoms in each molecule of the feedstock. Likewise, the cost to sequester a given mass of CO_2_,

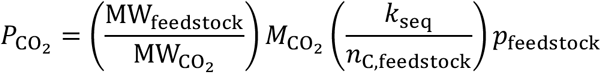

Where *p*_feedstock_ is the price per unit mass of the feedstock.

In this work, we succeeded in reducing *k*_seq_ from 525 to 1.0 (almost 3 orders of magnitude) (**Figure 8**). With 1% dunite and 24 hours of leaching, the approximate *k*_seq_ is 525 carbon invested/carbon sequestered. One method to reduce *k*_seq_ is to increase the time of leaching. After 96 hours of leaching, the magnesium concentration increased by 81% when compared to 24 hour leaching at the same pulp density (**Figure 3**). When increasing the time of leaching to 10 days, the magnesium concentration improved by an average of 147% (**Figure 8**). Increasing the pulp density of dunite also reduces *k*_seq_. When compared to 1% pulp density of dunite, increasing to 10% pulp density resulted in a 5-fold increase in magnesium leaching after 24 hours (**Figure 8**). When combining 10% pulp density with 10 days of leaching, magnesium leaching increased by 9.2 times that of 1% pulp density leached for 1 day (**Figure 8**), resulting in a *k*_seq_ of 57, since glucose input remained unchanged. For the lowest *k*_seq_ values achieved, fine-particle dunite at a pulp density of 60% was leached for 19 days with three different methods (**Figure 8** Leaching Attempts #12-14). The first method (**Figure 8** Leaching Attempt #12) was direct addition of 60% dunite, the second method (Leaching Attempt #13) was a 2-step dunite addition, where 30% dunite was added on day 0 and another 30% added on day 10, and the third method (Leaching Attempt #14) was a 2-step dunite addition where the first 30% dunite feed was removed from the leachate before the second 30% addition on day 10. All three leaching methods resulted in a *k*_seq_ equal to or less than 1.0, with the lowest *k*_seq_ being 0.96 on Leaching Attempt #13 in **Figure 8**.

**Figure 8.**
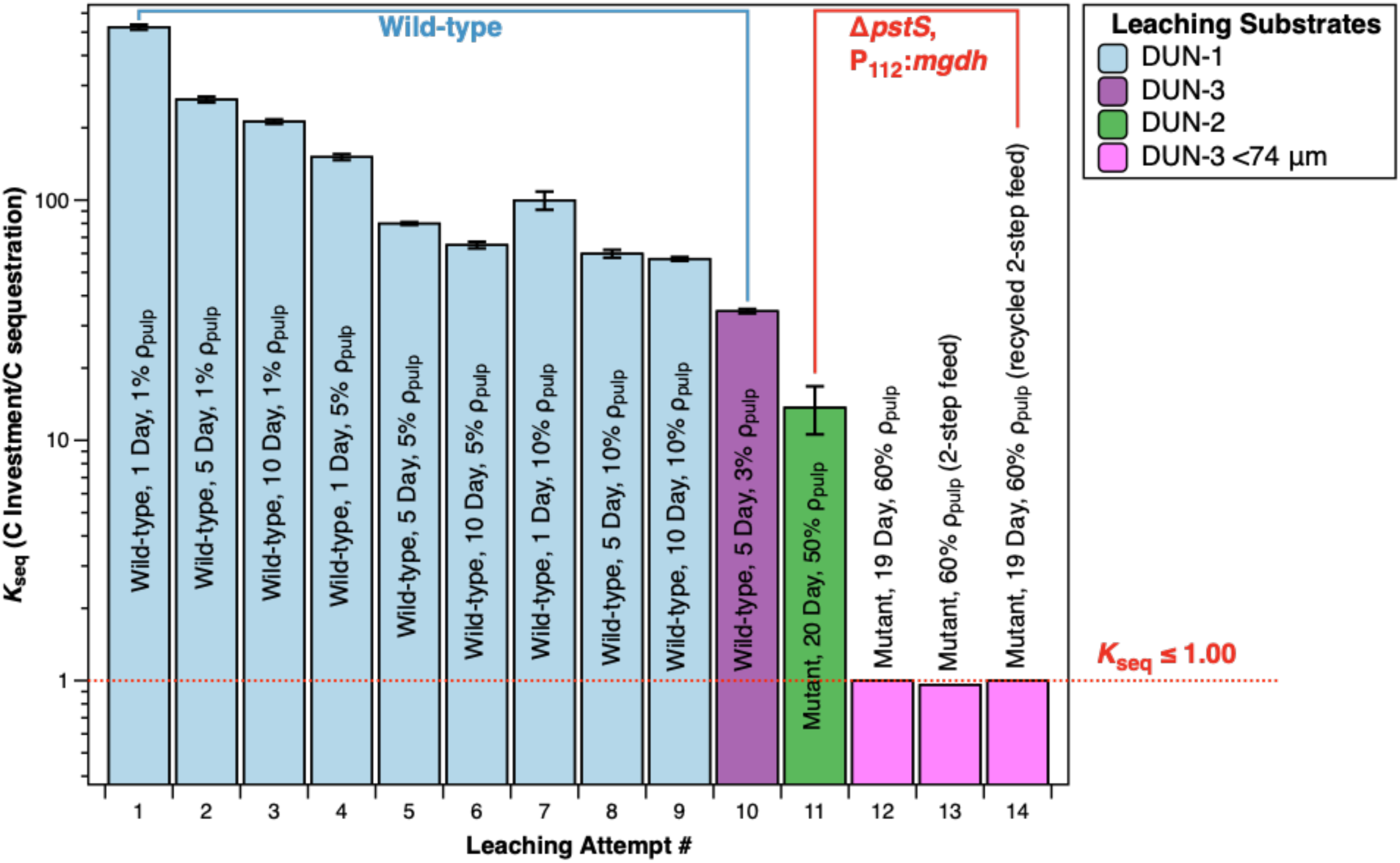
Process and genetic improvements reduce the amount of feedstock carbon needed to potentially sequester one CO_2_ molecule from 525 to 1. Increasing leaching time, pulp density, and reducing average particle sizes were able to reduce *k_seq_* (glucose carbon investment vs. carbon sequestration ratio) by several orders of magnitude. Increasing dunite pulp density from 1% to 10% had a larger impact in reducing *k*_seq_ than increasing leaching time from 1 to 10 days. Leaching studies with 60% pulp density with the Δ*pstS*, P_112_:*mdgh* double mutant over 19 days was able to reduce *k*_seq_ to 1.0 in three separate experiments (Leaching Attempts #12-14).

What implications do these results have for CO_2_ sequestration? For *k*_seq_ = 525, when using glucose as a feedstock, the cost of sequestering 1 tonne of CO_2_ is $358,000, a long way above the US Department of Energy’s (DOE) target of $100 per tonne^70^, and well above existing technologies^31^. On the other hand, if we could reduce *k*_seq_ to 1 (as in **Figure 8**) at industrial scale, the cost of carbon sequestration (at least the microbial feedstock part of the cost) through bio-accelerated weathering starts to become comparable to (still high) current technologies, at $682 per tonne^31^. If we could reduce *k*_seq_ to 0.15 (about a 6-fold reduction over our best attempt, much smaller than the reduction we achieved in the course of writing this article) then we could reduce the feedstock cost for sequestering 1 tonne of CO_2_ to $100. This would not account for the costs of mining ultramafic material, but nor does it account for the offset cost of energy-critical metal recovery. However, using this to sequester 20 gigatonnes of CO_2_ a year^5^ would require 2 billion tonnes of glucose, over 10-times current global annual sugar production^71^.

Sugars derived from cellulosic biomass are potentially much more abundant than glucose that is derived from cereal crops like corn, sugar cane, and sugar beet^72^. While there is no widespread production of cellulosic sugars today that is not immediately diverted to cellulosic ethanol production, this could happen in the future. Furthermore, it is estimated that cellulosic sugar production could be as low as $100 per tonne^73^ (in contrast to around $1,000 per tonne for glucose). Although cellulosic sugars are abundant, they are not limitless. Slade *et al*.^74^, made estimates of global bioenergy supply, and concluded that several tipping points would occur as the amount of biomass withdrawn from the biosphere for bioenergy (or some other chemical production, such as carbon sequestration) increased, forcing increasingly unpalatable choices on global society including mass adoption of vegetarian diets, mass deforestation, population control, or combinations thereof. Their analysis suggests that the first global agricultural tipping point (choosing one of the three unpalatable options) would occur when biomass was withdrawn at a rate of around 7 gigatonnes per year^61^. Assuming that cellulosic sugars are on average a 50:50 mix of C_5_ and C_6_ sugars, then to reduce the cellulosic biomass demand to sequester 20 gigatonnes of CO_2_ to 3.5 gigatonnes *k*_seq_ needs only to be reduced to 0.25.

### Possibilities for Downstream Carbon Mineralization

On its own, *G. oxydans* bioleachate saturated with magnesium ions will be difficult to convert into stable carbonate minerals. Due to the nature of acid leaching, the final pH of biolixiviants post-leaching tended to be low (4-5) in our experiments, making carbon mineralization chemically unfavorable. Since pH has a significant influence on the precipitation of carbonates due to the lower availability of carbonate ions in acidic conditions^75,76^, pH adjustments are required to increase carbonate mineral yields downstream. Increasing the concentration of gaseous CO_2_ injected or sparged through the leachate increases the total dissolved inorganic carbon (DIC); however, this reduces the pH of the leachate, reducing carbonate ion availability and decreasing the saturation index of carbonate minerals^2,75^.

A possible downstream pathway to adjust the pH of leachates to make carbon dioxide complexation chemically favorable is to add ammonia from biological or natural sources to synthesize magnesite, MgCO_3_, or nesquehonite, MgCO_3_·3H_2_O. Adding ammonia to CO_2_-sparged magnesium chloride solutions at ambient temperatures was shown to completely convert free magnesium ions into nesquehonite in about ten minutes^77^. Furthermore, ammonia can be recycled in this process through recovery of ammonia using activated carbon, which has been shown to recover ammonium chloride salts and separate them into ammonia and hydrochloric acid at near ambient conditions^78^. Another possibility is to simply add sodium hydroxide that has been electrically synthesized from carbon-neutral pathways.

Microbes that accelerate carbon mineralization, such as cyanobacteria, have been observed to accelerate mineralization in legacy alkaline mine tailings^34,35^. Coupling bio-accelerated weathering from *Gluconobacter oxydans* with biotic downstream processes could maximize carbonate precipitation in a low-resource and energy efficient process that could be genetically engineered further.

## Conclusion

This article shows that *G. oxydans* possesses the capability to bioleach metals for energy technologies and carbon mineralization from dunite. At low pulp densities (1%), the *G. oxydans* organic acid based-biolixiviant is 20× more effective at Mg leaching than deionized water. At high pulp densities (60%), the biolixiviant is 3.2× better than even gluconic acid alone. This suggests that microbially-accelerated weathering of ultramafic materials could be an environmentally-friendly method for supplying Mg for mineralization of CO_2_, and metals like Ni and Co for sustainable energy technologies at rates that can far exceed equivalent abiotic methods.

Several pH swing methods have been shown to be highly effective for greatly accelerating olivine carbonation^7,79^. However, scaling up these technologies to meet the world’s CO_2_ removal requirements may involve high energy costs^80^ and environmental drawbacks like those found in strong acid leaching involved with large scale energy-critical metal mining. Other methods of accelerated weathering involving energy-intensive preprocessing of mineral feedstocks by crushing, grinding, and chemically treating ultramafic rocks also come with their own environmental and scaling drawbacks. *In situ* accelerated ultramafic weathering can result in environmental leaching and environmental drainage of harmful doses of nickel, chromium, and cobalt^59,81–83^. These methods also involve slow kinetics that depend on high energy inputs to be accelerated and enormous land use^7,28,80,84^.

Bio-accelerated weathering may be an alternative approach to mineral weathering for carbon mineralization. When engineered further, this strategy may prove an economically-viable and scalable option that avoids the environmental concerns with *in situ* mechanically-accelerated weathering and strong acid leaching^51,59,82,83,85^, as well as the energy costs involved in high temperature and pressure reactions^15,80^. Energy requirements for heating in common abiotic carbonation processes, such as ammonium salts-driven pH swings and HCl pH swings, often result in CO_2_ emissions greater than the amount sequestered^80^. *G. oxydans*-based bioleaching was shown to be effective at ambient temperature and pressure, removing a major energy burden for this process.

In this article, we were able to reduce the number of carbon atoms in glucose needed to sequester one molecule of CO_2_ from 525 to 1. However, further work needs to be done to optimize *G. oxydans* for enhanced weathering for carbon mineralization. Maximizing the leaching of magnesium while minimizing carbon inputs and lowering the economic costs are key steps to making this technology viable for eliminating the several gigatonnes of carbon dioxide in excess on our planet.

## End Notes

### Data Availability

Datasets generated and analyzed during this study are included as **Dataset S1,** data for plots are deposited on GitHub at https://github.com/barstowlab/article-037-ultramafic-bioleaching and are archived on Zenodo^86^.

### Code Availability

No novel code was generated for this article.

### Materials & Correspondence

Correspondence and material requests should be addressed to B.B. and E.G..

### Author Contributions

Conceptualization, J.J.L., E.G., G.G., and B.B.; Methodology, J.J.L., S. Medin, S. Marecos., L.P, J.K., E.G., M.C.R., and B.B.; Investigation, J.J.L, L.P., and B.P.; Writing—Original draft, J.J.L.; Writing—Review and editing, J.J.L. and B.B.; Funding acquisition, B.B., G.G., and E.G.; Resources, M.C.R., E.G., and B.B.; Supervision, B.B. and E.G; Data curation, J.J.L. and B.B.; Visualization, J.J.L.; Formal analysis, J.J.L..

## Acknowledgements

This work was supported by Cornell University startup funds, a SPROUT award from the Cornell University College of Engineering, a Career Award at the Scientific Interface from the Burroughs-Welcome Fund to B.B., a gift from Mary Fernando Conrad and Tony Conrad to B.B., and gift from Nancy and Bob Selander to B.B. and E.G., and the U.S. Army Advanced Civil Schooling Program. J.J.L. was supported by a Cornell University graduate fellowship. S. Marecos was supported by a Link Foundation Graduate Fellowship. Daniel Schell at the National Renewable Energy Laboratory provided a sample of cellulosic hydrolysate. Lyndsey Fisher in the Department of Earth and Atmospheric Sciences at Cornell University processed ultramafic samples.

## Materials and Methods

### Substrates for Leaching

Ultramafic samples were collected from the Day Brook Dunite (35° 58’ 0.45” N, 82° 16’ 58.60” W) in western North Carolina^87^. The chemically weathered exterior of the dunite was removed using a tile saw (151991, MK Diamond Products, Inc., Torrance, CA). The trimmed interior was crushed into powder using a SPEX SamplePrep 8000M mill and a tungsten carbide vial. Before bioleaching experiments, the mineralogy of powder from two samples was analyzed via x-ray diffraction (XRD) with a Bruker D8 Advance ECO powder diffractometer at Cornell Center for Materials Research (CCMR). The instrument is equipped with a Copper K-α X-ray source. Analyses were made at 5 to 60 degrees, with an increment of 0.019 degrees and a dwell time of 0.3 seconds per step. To mitigate Fe fluorescence the detector’s lower discriminator was raised from 0.110 V to 0.182 V. XRD shows that sample DUN-1 consists of 64 wt. % olivine, 27 wt. % lizardite, 5 wt. % antigorite, and 4 wt. % talc. Sample DUN-2 contains 37 wt. % olivine, 59 wt. % lizardite, 2 wt. % talc, and 2 wt. % meixnerite. Sample DUN-3 contains 92 wt. % olivine, 5 wt. % antigorite, 2 wt. % lizardite, and 1 wt. % talc. Major and trace element compositions of the bulk rock were determined at the Hamilton Analytical Laboratory in Clinton, NY via XRF and LA-ICPMS and are shown in **Tables S1** and **S2**.

Three separate dunite samples were used in bioleaching studies. DUN-1 was used in the leaching studies shown in **Figures 3, 4, 5, 6**, and **Supplementary Figures S2 and S3**. Due to the depletion of DUN-1 in previous experiments, DUN-2 was used to produce the data used in **Figure 7**. DUN-3 was used in Leaching Attempts #10 and #12-14 in **Figure 8**. DUN-1 was used for all other leaching experiments.

### Microorganisms and Media

*Gluconobacter oxydans* strain NRRL B-58 (*Go*B58) was obtained from the American Type Culture Collection (ATTC), Manassas, VA. In all experiments, *Go*B58 was cultured in Yeast-Peptone-Mannitol media (YPM; 5 g L^−1^ yeast extract (C7341, Hardy Diagnostics, Santa Maria, CA), 3 g L^−1^ peptone (211677, BD, Franklin Lakes, NJ), 25 g L^−1^ mannitol (BDH9248, VWR Chemicals, Radnor, PA)).

Gluconobacter oxydans mutant strain Δ*pstS*, P_112_:*mdgh* contains a clean deletion of the gene encoding the periplasmic phosphate-binding protein, PstS, and an up-regulation of the membrane-bound glucose dehydrogenase gene with promoter P_112_. This mutant strain was previously constructed by Schmitz *et al.*^43^.

### Growth and Organic Acid Production

*Go*B58 was inoculated into YPM liquid culture then incubated for 24 hours at 30°C and 200 rpm in a shaking incubator (Infors HT Multitron). Cultures were then back-diluted to OD590 of 0.05 in a total volume of 25 mL YPM in 50 mL flasks. The flasks were then incubated at 30°C and 200 rpm until the OD590 reaches approximately 1.5 after about 48 hours. Cultures were then split into 7.5 mL triplicates with 7.5 mL of 40% glucose added to each triplicate, for a total volume of 15 mL. The glucose and bacterial mixture was then incubated for 48 hours at 30°C and 200 rpm, starting biolixiviant production. After 48 hours, the pH of each flask is measured (VWR Symphony B10P.).

Cellulosic hydrolysate (National Renewable Energy Laboratory, Golden, CO) was used in some leaching studies as an alternative glucose feedstock. In experiments using cellulosic hydrolysate, it was diluted to 80% concentration so that the total sugar concentrations (**Table S2**) would be equal to 40%. Then 7.5 mL of 80% cellulosic hydrolysate was added in replacement of 7.5 mL of 40% glucose.

### Bioleaching Studies

Bioleaching experiments were conducted with dunite at a pulp density of 1-10% wt/vol, as specified. Minerals were added to 15 mL of biolixiviant, then agitated at room-temperature in a shaking incubator for 24 hours. After 24 hours, the ultramafic rock and biolixiviant slurry was centrifuged and the bioleachate was extracted for ICP-MS analysis.

### Analytical Methods

Leachates from bioleaching experiments were filtered through a 0.45 µm AcroPrep Advance 96-well filter plate (8029, Pall Corporation, Show Low, AZ, USA) by centrifugation at 1,500 × g for 5 min, then diluted to 1% with 2% trace metal grade nitric acid (JT9368, J.T. Baker, Radnor, Pennsylvania, USA). Samples were analyzed with an Agilent 7800 ICP-MS. The concentrations of metals were measured against a custom ICP-MS standard solution containing 10 mg/L of Al, As, B, Ca, Cd, Co, Cr, Cu, Fe, K, Mg, Mn, Mo, Na, Ni, P, Pb, S, Sb, Se, Si, Sn, and Zn (High Purity Standards, North Charleston, SC) and an ICP-MS internal standard (5188-6525, Agilent Technologies, Santa Clara, CA). Quality checks were done with intermittent standards and blanks as specified in the ICP-MS guidelines and protocols^88^.

### Statistics

A Welch’s two sample *t*-test was performed to determine statistical significance in comparative bioleaching studies. A Welch’s two sample *t*-test was done since some studies, such as a comparison of sugar feedstocks (**Figure 5**), resulted in differences in variances between the two groups tested. The Welch’s *t*-test is more appropriate in instances when the variances between two groups cannot be considered as equal. Significance was displayed in all figures as dots and stars, where *p* < 0.1 (.), *p* < 0.05 (*), *p* < 0.01 (**), *p* < 0.001 (***).

