## Supplementary Information for "Bio-Accelerated Weathering of Ultramafic Minerals with *Gluconobacter oxydans*"

#### Supplementary Information Figures

**Figure S1.** Dunite metal composition.

**Figure S2.** Aluminum and chromium leaching results.

**Figure S3.** Stoppered vs. unstoppered bioleaching results.

#### Supplementary Information Tables

**Table S1.** Dunite metal oxide composition.

**Table S2.** Dunite trace element composition.

**Table S3.** Cellulosic hydrolysate composition.

**Table S4.** Metal concentrations in cellulosic hydrolysate.

#### Supplementary Information Datasets

**Dataset S1.** Raw ICP-MS measurements for **Figures 2 8, Figure S3, and Table S4**.

#### Supplementary Information Notes

**Note S1.** Chemical reactions for indirect carbonation of forsterite.

#### Supplementary Information References

#### Supplementary Figures

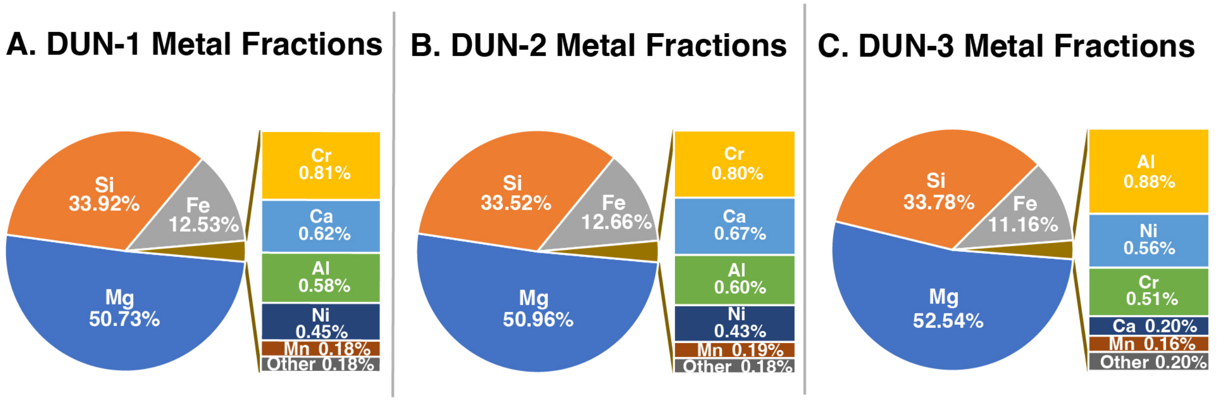


**Figure S1. Dunite metal fractions of two dunite samples used in bioleaching studies, DUN-1 (A) and DUN-2 (B).** Mg, Si, and Fe are the main metals in the dunite used in the experiments. This figure shows the fractions of pure metals contained in the dunite, excluding the molecular weight of the anionic oxide. Mineral compositions of both dunite samples are shown in **Table S1**. DUN-2 was used in leaching experiments shown in **Figure 4** and Leaching Attempt #11 in **Figure 8**. DUN-3 was used in Leaching Attempts #10 and #12-14 in **Figure 8**. DUN-1 was used for all other leaching experiments.

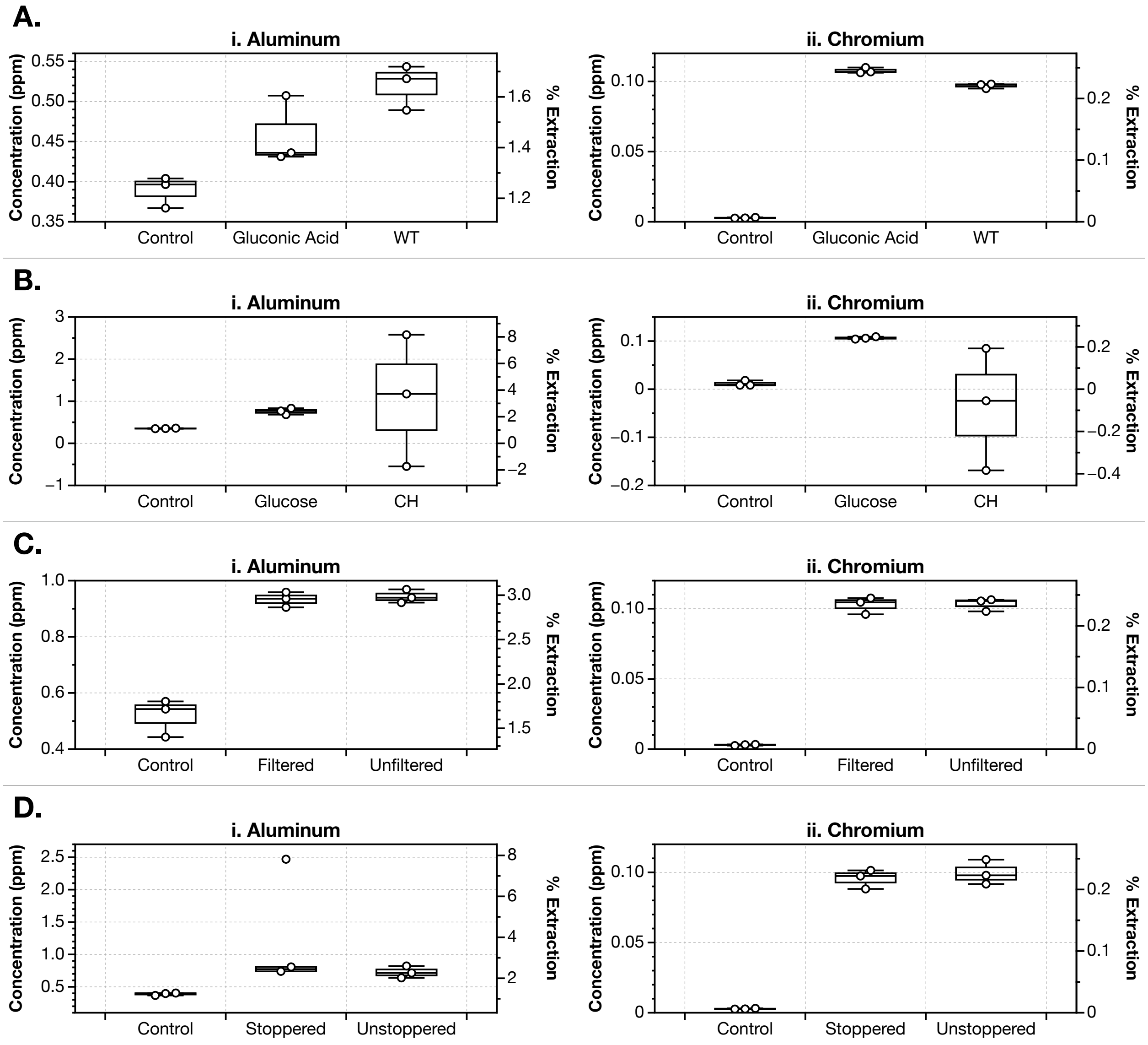


**Figure S2. Aluminum and chromium were excluded from main text figures and have lower leaching efficiencies compared to Mg, Mn, Fe, Co, and Ni.** The concentrations and extraction efficiencies of Al and Cr are plotted in bioleaching studies with 1% dunite and 24 hours of leaching. **(A)** *G. oxydans* leaching is compared to gluconic acid diluted to the same pH and deionized water (control). **(B)** Cellulosic hydrolysate-based biolixiviants are compared to glucose-based biolixiviants and a deionized water control. (**C)** Filtered biolixiviants are compared with unfiltered, whole-cell biolixiviants. **(D)** Stoppered leaching is compared with unstoppered leaching.

The relatively low extraction efficiency of Cr could be attributed to the fact that Cr is mainly contained in chromite mineral grains in the dunite samples. Chromite ($FeCr_2O_4$) is a highly stable mineral which is difficult to leach chemically or with acids [Mena1990a].

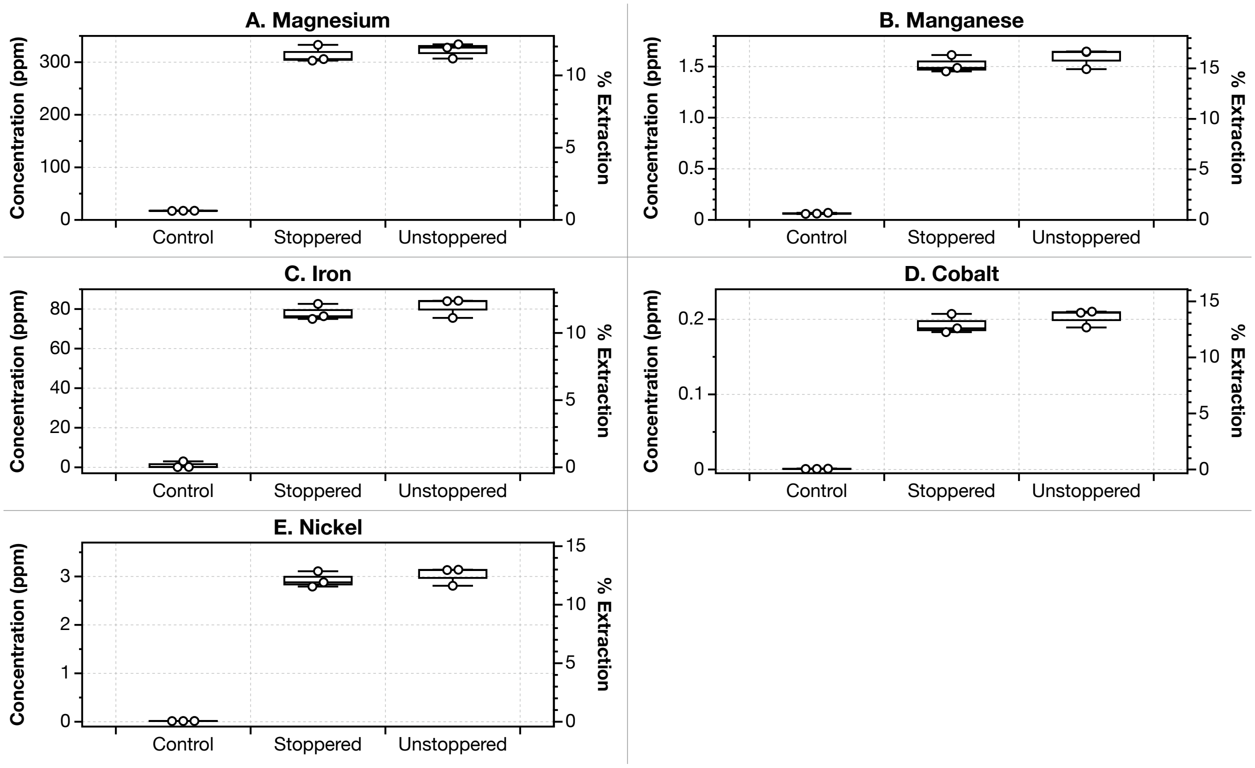


**Figure S3. Stoppering does not significantly affect bioleaching in a 24-hour leaching study.** To investigate the effect of using stoppers during bioleaching to prevent gas exchange, stoppered bioleaching (col. 2) was compared to unstoppered bioleaching (col. 3) with 1% dunite for 24 hours using wild-type *G. oxydans* biolixiviants. Unstoppered leaching resulted in an average leaching increase of 4% across all metals. However, this increase could be attributed to liquid volume loss due to evaporation in unstoppered leaching experiments, resulting in higher final metal concentrations. A Welch's two-sample t-test reveals no significant difference between both groups, with $p$-values between 0.3 and 0.6.

#### Supplementary Tables

| Oxide | Weight % DUN-1 | DUN-2 | DUN-3 |
|---|---|---|---|
| MgO | 45.58 | 46.07 | 48.46 |
| $SiO_2$ | 39.32 | 39.09 | 40.20 |
| FeO | 8.74 | 8.88 | 7.99 |
| $Al_2O_3$ | 0.60 | 0.62 | 0.93 |
| CaO | 0.47 | 0.51 | 0.16 |
| MnO | 0.13 | 0.13 | 0.12 |
| $Na_2O$ | 0.069 | 0.066 | 0.028 |
| $TiO_2$ | 0.043 | 0.043 | 0.045 |
| $K_2O$ | 0.019 | 0.020 | 0.043 |
| $P_2O_5$ | 0.014 | 0.014 | 0.060 |
| $Co_2O_3$* | 0.021 | - | - |
| Loss on ignition (LOI) | 3.76 | 3.09 | 0.76 |
| **sumMaj+LOI** | 98.74 | 98.54 | 98.80 |
| **sumAll** | 99.51 | 99.51 | 99.50 |

*Note: $Co_2O_3$ was measured in a separate XRF analysis of DUN-1 and was appended to this table

**Table S1. Metal oxide composition of dunite samples as weight percentages, in order of abundance.** DUN-1 (col. 1) was used in the majority of bioleaching experiments, and DUN-2 (col. 2) was only used for the experiment shown in **Figure S2** in the main text due to the depletion of DUN-1. Ni and Cr concentrations are displayed as trace elements in **Table S2**. Row "sumMaj + LOI" refers to the sum of major oxides listed + Loss on ignition (LOI). Row "sumAll" is the sum of major oxides, LOI, volatile components, and trace elements detected.

| | Parts per million (ppm) | | |
|---|---|---|---|
| **Element** | **DUN-1** | **DUN-2** | **DUN-3** |
| Ni | 2,417 | 2,334 | 3,099 |
| Cr | 4,391 | 4,366 | 2,813 |
| V | 20.55 | 17.97 | 12.23 |
| Sc | 5.89 | 4.37 | 2.29 |
| Cu | 4.08 | 12.90 | n.d. |
| Zn | 54.09 | 54.92 | 70.58 |
| Ga | n.d. | n.d. | n.d. |
| Ba | 11.91 | 9.56 | 22.48 |
| Rb | n.d. | 0.77 | n.d. |
| Cs | 4.08 | 0.54 | 4.13 |
| Sr | 0.60 | 0.88 | 3.47 |
| Y | 1.99 | 3.37 | 2.33 |
| Zr | 11.70 | 10.63 | 14.69 |
| Hf | 0.16 | 0.68 | n.d. |
| Nb | n.d. | n.d. | 0.67 |
| Ta | 1.10 | 1.48 | 0.97 |
| Mo | n.d. | n.d. | 0.92 |
| La | 0.27 | n.d. | n.d. |
| Ce | 6.80 | 2.82 | 13.21 |
| Nd | 6.05 | 6.55 | 6.23 |
| Sm | 1.34 | 1.72 | 1.33 |
| Dy | 1.31 | 1.51 | 1.19 |
| Yb | 0.93 | 1.09 | 1.23 |
| Th | 1.17 | n.d. | n.d. |
| U | 1.35 | 2.45 | 1.03 |
| Tl | n.d. | n.d. | 0.05 |
| Pb | n.d. | n.d. | n.d. |
| Sn | 2.46 | 4.22 | 4.44 |
| Bi | n.d. | n.d. | n.d. |
| Sb | n.d. | n.d. | n.d. |

n.d. = not determined

**Table S2. Trace element composition of dunite samples DUN-1 and DUN-2.**
1,000 parts per million (ppm) = 0.1 weight %.

| **Sugars/Component** | **Concentration (mg/mL)** |
|---|---|

| | |
|---|---|
| Glucose | 311.97 |
| Xylose | 170.03 |
| Galactose | 1.66 |
| Arabinose | 17.10 |
| Frutose | 0.00 |
| Lactic Acid | 1.24 |
| Glycerol | 0.00 |
| Acetic Acid | 1.24 |
| Ethanol | 0.00 |
| HMF | 0.00 |
| Furfural | 0.00 |

**Table S3. Concentrations of sugars and other components in cellulosic hydrolysate sample.**

| Element | Concentration (mg/L) | Standard Deviation |
|---|---|---|
| Mg | 1321.13 | 56.48 |
| Al | 75.12 | 3.82 |
| Cr | 5.33 | 0.31 |
| Mn | 29.74 | 1.73 |
| Fe | 125.07 | 31.21 |
| Co | 0.38 | 0.02 |
| Ni | 11.02 | 0.61 |
| Cu | 0.23 | 0.01 |
| Zn | 8.55 | 0.65 |

**Table S4. Concentrations of metals in cellulosic hydrolysate sample measured by ICP-MS.**

### Supplementary Notes

#### Note S1. Chemical Reactions for Indirect Carbonation of Forsterite

$Mg_2SiO_{4(s)} + 4C_6H_{12}O_{7(aq)} \rightarrow 2Mg^{2+}_{(aq)} + 4C_6H_{11}O_7{}^{-}{}_{(aq)} + 2H_2O + SiO_{2\,(s)}$ **(S1)**

$CO_{2(g)} + H_2O \rightarrow H^{+}_{(aq)} + HCO_3{}^{-}{}_{(aq)}$ **(S2)**

$Mg^{2+}_{(aq)} + HCO_3{}^{-}{}_{(aq)} \rightarrow MgCO_{3(s)} + H^{+}_{(aq)}$ **(S3)**

Indirect carbonation, where $CO_2$ injection occurs as a separate step after magnesium leaching, has a higher conversion rate of carbon dioxide to carbonates and has faster kinetics when compared to direct carbonation [Pan2015a]. In indirect carbonation, an extractant is used to free magnesium ions from magnesium silicates before carbonation (**Eqn. S1**). This precipitates silica, where it can be removed from the aqueous solution through a filter [Pan2015a, OConnor2005a, Turri2019a]. Stirring is required throughout dissolution to prevent accumulation of silica on forsterite surfaces that can inhibit dissolution [Béarat2006a, Turri2019a]. By removing silica from the aqueous solution before carbonation, the remaining solution is rich in magnesium ions, improving the reactive favorability and kinetics of carbon mineralization [Addassi2024a]. Carbon dioxide is then dissolved into the solution (**Eqn. S2**), where it can then react with magnesium ions to precipitate magnesite (**Eqn. S3**). In this process, each magnesium ion liberated from forsterite has the storage capacity for one carbon dioxide molecule. **Eqns. S1-S3** show the catalyzed reaction when using gluconic acid as an extractant on forsterite, $Mg_2SiO_4$, the magnesium endmember of olivine. A schematic of a proposed system for bio-accelerated weathering coupled with downstream carbonation and metal recovery is shown in **Figure 1** of the main text.
